# Action-related brain dynamics during leader–follower coordination

**DOI:** 10.64898/2026.10.11.758328

**Authors:** Santeri Ruuskanen, Juan Camilo Avendano-Diaz, Riitta Hari, Lauri Parkkonen

## Abstract

Leader–follower dynamics are essential for social interaction, enabling smooth inter-personal coordination. Although neuroimaging studies have identified brain networks associated with leading and following, the dynamical brain signatures supporting these roles remain poorly characterized. Here, sixteen dyads performed a joint-improvisation mirror-game task with predefined leader–follower roles while magnetoencephalography (MEG) and hand kinematics (acceleration) were simultaneously recorded. We investigated role-specific modulations of source-space MEG activity in delta (0.5–4 Hz), theta (4–8 Hz), alpha (8–13 Hz), beta (13–30 Hz), and gamma (40–90 Hz) frequency bands, as well as coherence between cortical activity and movement kinematics. Compared with leaders, followers exhibited stronger theta-band activity in a cluster comprising frontal, cingulate, and sensorimotor regions, and stronger coherence between movement kinematics and occipital cortical activity, consistent with enhanced monitoring, adaptation, and reliance on visual kinematic information. In contrast, leaders showed reduced alpha-band power in sensorimotor and premotor regions, and stronger beta-band activity in occipital cortices, likely reflecting greater demands for action planning and predictive control. Our results identify complementary spectral and action-related brain signatures associated with leader–follower coordination during joint improvisation, providing new insights into the neural underpinnings of real-time interpersonal coordination.

## Introduction

Human social interaction depends on the ability of individuals to smoothly coordinate their actions in time and space. Such coordination spans a wide range of behavior, from conversational turn-taking to complex joint actions, such as music and dance performances (Sebanz et al. 2006; Keller et al. 2014; Foster Vander Elst et al. 2023). In these contexts, coordination unfolds continuously and requires ongoing flexible adaptation to the actions of the partners (Bolt and Loehr 2021; Foster Vander Elst et al. 2023).

Research on the brain basis of social interaction has increasingly adopted more naturalistic experimental setups. This shift has been motivated by calls to study social interaction as it naturally occurs between interacting individuals, giving rise to two-person and second-person neuroscience approaches that enable the investigation of reciprocal and dynamic processes during real-time interaction (Hari and Kujala 2009; Schilbach et al. 2013; Hari et al. 2015).

During efficient coordination, interacting individuals often adopt complementary roles characterized by unequal levels of adaptation, commonly described as leader–follower dynamics (Fairhurst et al. 2014; Keller et al. 2014; Chauvigné and Brown 2018; Chauvigné et al. 2018; Vanzella et al. 2019; Bolt and Loehr 2021). In such interactions, one partner prioritizes maintaining an internally generated action plan (e.g., tempo or movement structure), while the other allocates greater resources to monitoring and adapting to the partner’s behavior to sustain coordinated action (Fairhurst et al. 2014, Konvalinka et al. 2014). Leader–follower roles can be explicitly assigned (e.g., in ensemble music or couple dancing) or they can emerge dynamically during interaction (e.g., conversational turn-taking, cooperative object manipulation, or joint improvisation) (Keller et al. 2014; Chauvigné et al. 2018; Bolt and Loehr 2021; Foster Vander Elst et al. 2023).

Neuroimaging studies have begun to characterize the neural correlates underlying leader–follower dynamics. Using functional magnetic resonance imaging (fMRI) in rhythmic finger-tapping (Fairhurst et al. 2014) and bimanual coordination tasks (Chauvigné et al. 2018; Chauvigné and Brown 2018), the leader role has been associated with greater recruitment of action-related networks involved in internally driven action control, prospective planning, and self-monitoring. In contrast, the follower role has been associated with stronger engagement of sensory and social–cognitive networks that support adaptation to external actions (Chauvigné et al. 2018). Moreover, task-related activity in these networks has been shown to correlate with expertise in the respective roles (Chauvigné and Brown 2018). Together, these findings highlight the complementary nature of leader and follower roles, with leaders relying more strongly on internally generated action control and followers allocating greater resources to externally driven processes and continuous adaptation (for a review, see Bolt and Loehr 2021).

While these studies have identified spatially distinct brain activation patterns associated with the leader and follower roles, the brain’s oscillatory dynamics linked to these coordination patterns remain less characterized. Evidence from EEG hyperscanning during a joint finger-tapping task suggests that the level of alpha-band activity may differentiate leaders from followers, with spontaneously emerging leaders showing stronger alpha suppression in frontal regions during joint action (when the recordings were referenced to the nose), which was interpreted as increased allocation of resources to self-related processing and prospective control (Konvalinka et al. 2014). MEG hyperscanning during auditory communication in a musical turn-taking task further revealed stronger occipital alpha- and parieto-occipital beta-band activity in followers than in leaders, interpreted as reflecting increased working-memory and attentional demands associated with the follower role (Yoneta et al. 2022). However, these effects were observed during preparatory imagery rather than ongoing coordination. Beyond role differentiation, alpha oscillations have been shown to covary with the balance between self–other integration and segregation in joint action (Novembre et al. 2016), whereas beta-band activity has been related to the maintenance of interpersonal coordination (Novembre et al. 2017). Nonetheless, it remains unclear how brain oscillatory dynamics support leader–follower roles during ongoing, reciprocal interpersonal coordination.

The coupling between brain activity and motor action during leader–follower interactions may provide critical insight into the coordination dynamics and self–other integration processes supporting the respective roles. Visual frequency-tagging studies have shown that early visual regions differentially track self- and other-generated movements, with followers exhibiting stronger neural responses to the partner’s movements and leaders showing enhanced tracking of self-generated motion (Varlet et al. 2020). Follow-up work further revealed stronger neural tracking of visual self–other synchronization errors in followers, associated with greater adaptation demands, and earlier, shorter-latency tracking in leaders, potentially reflecting top-down predictive processes (Varlet et al. 2025). Complementary evidence from interactive sensorimotor setups suggests that distinct cortical systems track self- and other-related movement kinematics beyond visual processing (Dumas et al. 2020) and that the kinematic phase of hand movements modulates occipital beta-band activity more strongly in followers than in leaders, potentially reflecting distinct strategies for integrating visual kinematic information (Zhou et al. 2016). It remains an open question whether direct, frequency-specific brain–action coupling is modulated by leader–follower roles.

A direct and particularly suitable approach to quantifying continuous brain–action coupling during interpersonal coordination is provided by corticokinematic coherence (CKC). CKC indexes frequency-specific coupling between cortical activity and movement kinematics during rhythmic actions (Jerbi et al. 2007; Bourguignon et al. 2011, 2015; Piitulainen et al. 2013) and tracks both executed and observed actions (Bourguignon et al. 2011, 2013; Piitulainen et al. 2013; Marty et al. 2015).

Here, we employed our MEG hyperscanning setup (Zhdanov et al. 2015) in combination with spectral and corticokinematic coherence analyses to investigate the neural–behavioral underpinnings of leader–follower coordination during a continuous finger-movement mirror game with predefined interaction roles. The mirror game provides a controlled yet naturalistic setting for studying joint improvisation and interpersonal coordination dynamics (Noy et al. 2011). We explored role-dependent differences in cortical oscillatory activity in the delta (0.5–4 Hz), theta (4–8 Hz), alpha (8–13 Hz), beta (13–30 Hz), and gamma (40–90 Hz) frequency bands, and examined differences in frequency-specific coupling between cortical activity and movement kinematics across leader vs. follower roles. We hypothesized that followers would exhibit stronger coupling between movement kinematics and activity in visual regions, reflecting their greater reliance on external information during coordination (Zhou et al. 2016), while leaders would exhibit stronger coupling between kinematics and activity in sensorimotor regions, reflecting a greater relative contribution of proprioceptive input (Bourguignon et al. 2015) to the cortex during self-generated movements.

## Materials and Methods

### Subjects

Sixteen dyads (12 same-sex, 4 mixed-sex; 32 subjects; 14 female, 18 male; mean ± SD age 28.8 ± 4.8 years, range 21–41 years) participated in the experiment. The participants were mainly students or researchers in the Helsinki and Brussels metropolitan areas. All participants were determined right-handed according to the Edinburgh handedness inventory (Oldfield 1971) and had normal or corrected-to-normal vision. The participants gave written informed consent. The study received ethical approval from the Ethics Committee of the Hospital District of Helsinki and Uusimaa and was conducted according to the Declaration of Helsinki. Due to unavailable anatomical MRI or accelerometer data for some participants, and a missing trial for one dyad, we present results from 20 subjects (7 female, 13 male; age 29.0 ± 4.9 years).

### Experimental setup

We employed our unique MEG hyperscanning setup (later referred to as MEG2MEG), which enables the synchronized MEG recording of two interacting participants located at two separate MEG laboratories (Figure 1A) (Zhdanov et al. 2015). During the experiment, one participant was seated inside a three-layer magnetically-shielded room (MSR) (Imedco AG, Hägendorf, Switzerland) at the MEG Core of Aalto NeuroImaging at Aalto University in Espoo, Finland (later referred to as Aalto) and the other within a lightweight MSR (MaxShield^TM^; Megin Oy, Espoo, Finland) at the ULB-Hôpital Erasme in Brussels, Belgium (later referred to as Erasme). The two MEG sites were equipped with microphones and video cameras and were connected across the internet to allow the participants to interact seamlessly via a custom-protocol audiovisual link with an end-to-end latency of approximately 130 ms, low enough to enable naturalistic motor coordination (Zhdanov et al. 2015). A live image of the participant at the other site was projected onto the standard MEG-compatible back-projection screen. To enable offline synchronization of the recordings, GPS receivers were employed at both experimental sites to synchronize the device clocks and insert GPS-derived timestamps into the recorded video, audio, and MEG data. Further technical details on the MEG2MEG setup are provided in our previous publication (Zhdanov et al. 2015).

**Figure 1.**
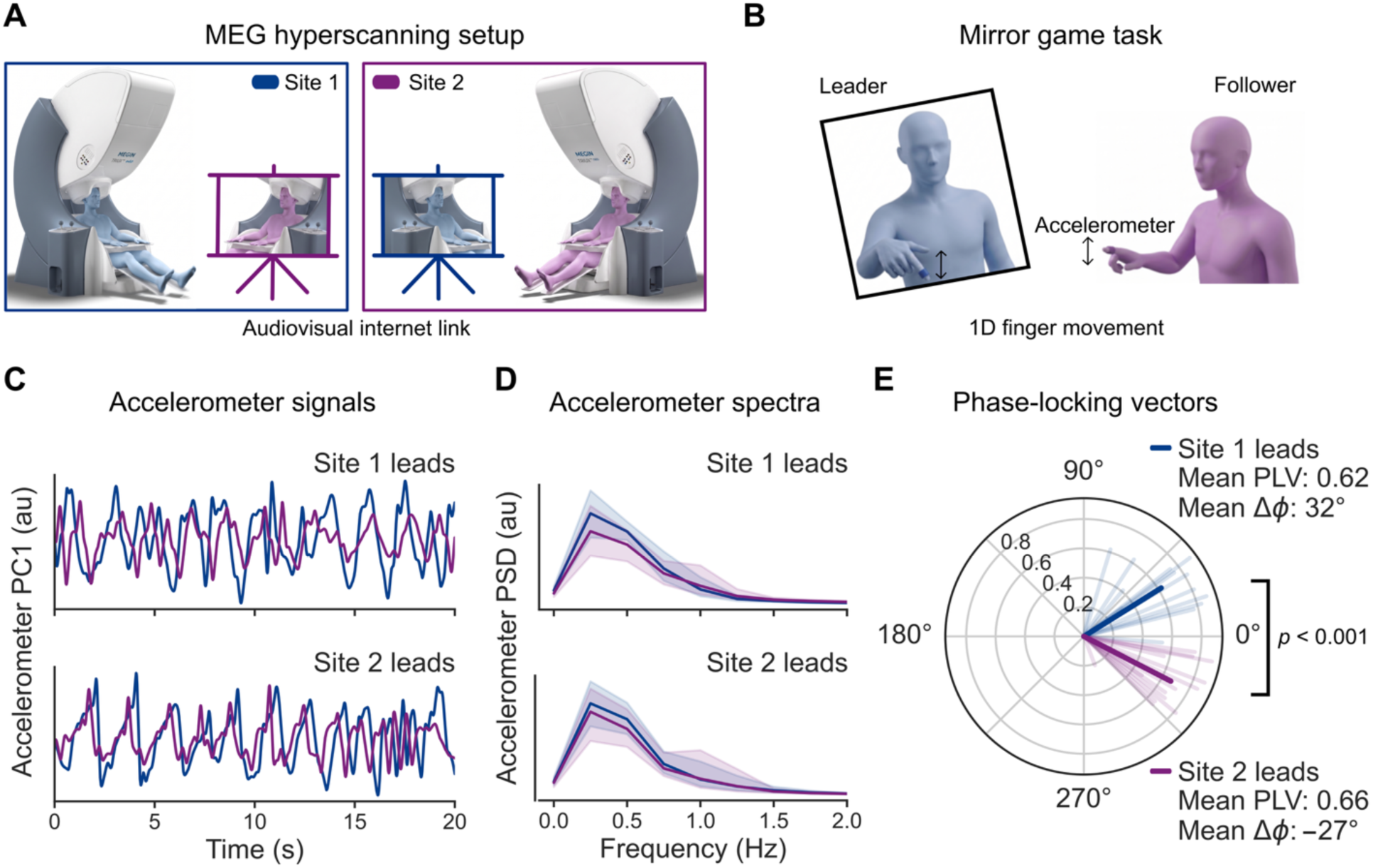
MEG hyperscanning setup, mirror-game task, and behavioral results. **(A)** MEG was recorded simultaneously from two interacting participants at different sites (Site 1 in blue, Site 2 in magenta) which were connected by a low-latency audiovisual internet link. **(B)** The participants performed a 1D finger-movement mirror-game task, moving their index finger vertically. **(C)** Accelerometer signals (first principal component; PC1) recorded from the index fingers of the participants exhibited clear behavioral synchrony. **(D)** Grand-average accelerometer spectra across subjects indicate variable movement frequency. The shaded areas correspond to the 95% bootstrap confidence interval of the mean. **(E)** Phase-locking vectors were computed separately from the blocks where participants at each site were assigned the leading role. The dark thick blue and magenta lines indicate the grand-average vectors. The light thin lines represent individual dyads.

### Behavioral task

The participants performed an adapted version of the mirror game (Noy et al. 2011). Here, each participant performed vertical hand movements with their pointed index finger dragging along a transparent touch plate positioned on a table in front of them (Figure 1B). The participants were instructed to perform the movement using only their wrists while keeping the rest of the arm immobile. They were encouraged to improvise novel movement patterns, varying the amplitude and speed of the movement freely, as long as the movement was continuous throughout each block. The experiment included seven conditions: rest, moving in isolation with the screen blank, moving while the other participant was observing, observing while the other participant was moving, synchronous joint improvisation with the other participant, leading, and following. We recorded three 60-s blocks of each condition in random order. In the current study, we only examine and discuss the leader–follower interaction. Leaders were instructed to improvise movements continuously without waiting for or responding to the follower, while ensuring that the movements remained feasible to imitate. Followers were instructed to reproduce the leader’s movements as closely and as synchronously as possible.

### Recordings

We recorded 306-channel MEG, vertical and horizontal electrooculogram (EOG), single-lead electrocardiogram (ECG), finger acceleration, audio, and video signals simultaneously from both participants of each dyad. MEG and physiological signals were recorded using two very similar whole-scalp MEG devices, namely Elekta Neuromag® (Megin Oy, Espoo, Finland) at Aalto and Vectorview^TM^ (Megin Oy, Espoo, Finland) at Erasme. The sensor arrays of both devices comprise 102 triplets of two perpendicularly oriented planar gradiometers and a single magnetometer. Bipolar EOG electrodes were placed on the temples to record horizontal eye movements, and on the zygomatic bone and the frontal bone to record vertical eye movements and eye blinks. Single-lead ECG was recorded between the clavicles. The finger movements were tracked using a 3-axis accelerometer (ACC; ADXL335 iMEMS Accelerometer; Analog Devices, Inc., Norwood, MA, USA) mounted on the index finger. These data were bandpass-filtered to 0.1–330 Hz and sampled at 1 kHz. The head position of the subject was continuously monitored using four head position indicator (HPI) coils at Erasme and five such coils at Aalto. The locations of the HPI coils relative to anatomical landmarks on the head were digitized using a Polhemus Fastrak electromagnetic motion tracker (Polhemus, Inc., Colchester, VT, USA). Audio and video data were recorded in the Helsinki VideoMEG format (Zhdanov et al. 2018) and were used during visual inspection of the MEG signals.

At Aalto, structural T1-weighted MRI data were recorded using a 3-T Siemens Magnetom Skyra (Siemens Healthineers AG, Erlangen, Germany) at the Advanced Magnetic Imaging Center of Aalto NeuroImaging (MPRAGE sequence; repetition time (TR) 2530 ms; echo time (TE) 3.3 ms; flip angle 7 deg; inversion time (TI) 1100 ms). At Erasme, 1.5-T Philips Intera and Philips Achieva (Koninklijke Philips N.V., Amsterdam, The Netherlands) were employed (T1 TFE IR sequence; TR ∼8.7 ms; TE ∼4.2 ms; flip angle 8 deg).

### Analysis of accelerometer data

The 3-axis accelerometer (ACC) signals were bandpass-filtered to 0.3–195 Hz using a finite impulse response (FIR) filter with the default parameters in MNE-Python (version 1.9) (Gramfort et al. 2014, 2013). Any remaining DC component was then removed by subtracting the channel-wise mean from the data for each 60-s block. The 3-axis data were then collapsed into a single time series utilizing the first principal component (PC1) for the behavioral analysis. This component corresponds to the direction of acceleration with the largest variance in the 3-dimensional space. For the corticokinematic coherence (CKC) analysis, we employed the Euclidean norm of the three channels, consistent with prior CKC studies (Bourguignon et al. 2011).

To estimate the power spectral density (PSD) of the ACC signals, we employed Welch’s method (Welch 1967) with 4-s epochs and 3.6-s overlap. To verify that the participants performed the task as instructed, we computed the phase-locking vectors (Lachaux et al. 1999) between the ACC signals in each dyad separately from the blocks where each site led the movement. To that end, the ACC PC1 signals *x*_1_(*t*), *x*_2_(*t*) were first bandpass-filtered to 0.1–3 Hz. Analytic signals were then obtained through the Hilbert transform *H*(⋅) as *z*_1_(*t*) = *H*(*x*_1_(*t*)), *z*_2_(*t*) = *H*(*x*_2_(*t*)) and concatenated across blocks in the time dimension. Phase-locking vectors and phase-locking values were then calculated separately for the leader and follower conditions for each subject as

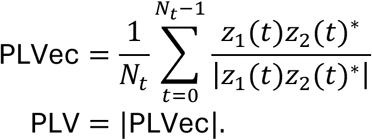

Lastly, PLVecs and PLVs were averaged separately for the condition in which the participant at Aalto (Site 1) was leading and the condition in which the participant at Erasme (Site 2) was leading.

### Preprocessing of MEG data

To ensure data quality and participant compliance, we inspected the MEG, physiological, and kinematic data together with synchronized video recordings using the locally developed and publicly available *mne-videobrowser* tool (https://github.com/ttaiv/mne-videobrowser). To suppress external interference and correct for head movement during the measurement, we applied spatiotemporal signal-space separation (tSSS; Taulu and Simola 2006) utilizing the implementation in MaxFilter software (Megin Oy, Espoo, Finland; version 2.2.30; correlation limit 0.9; correlation window length 20 s). The data were subsequently bandpass-filtered to 0.1–195 Hz. To suppress physiological artifacts due to cardiac activity, eye movements, and eye blinks, we employed Independent Component Analysis (ICA) by applying the MNE-Python implementation of the FastICA algorithm (Hyvärinen 1999). Independent components (ICs) were automatically pre-selected using Pearson correlation with the EOG channels and cross-trial phase statistics with the ECG channel. All ICs were then inspected visually, and the components containing clear artifacts were removed. The cleaned data from both participants in a dyad were synchronized utilizing GPS timestamps included at recording time. The data were then bandpass-filtered to 0.3–195 Hz and split into 4-s epochs with 3.6-s overlap. Finally, epochs with gradiometer signals containing zero-to-peak amplitudes greater than 700 fT/cm were rejected. Only gradiometer data were retained for further analysis.

### Processing of MRI data and MEG forward model

T1-weighted structural MRI volumes were automatically segmented to create individual 3-D cortical reconstructions for each participant using FreeSurfer software (Dale et al. 1999; Fischl et al. 1999). To model the individual head conductivity profile and solve the MEG forward problem, we employed a single-shell boundary element method (BEM) volume conductor model. The intracranial volume boundary was tessellated using the FreeSurfer watershed algorithm (Ségonne et al. 2004). We defined a cortically-constrained surface source space with recursively divided icosahedron spacing (ico4) for each participant, which resulted in 2562 source points for each cortical hemisphere. The anatomical data were coregistered with the MEG recordings using the MNE-Python coregistration GUI.

### Spectral analysis

To investigate how leader and follower roles modulate cortical oscillations, we estimated the source-level power-spectral density (PSD) in these roles using the Dynamic Imaging of Coherent Sources (DICS) method (Gross et al. 2001). We computed the cross-spectral density (CSD) matrices of the MEG data from the Fast Fourier Transform (FFT) coefficients of the overlapping epochs described above. Source-level PSD was then estimated using the MNE-Python DICS implementation in the source orientation that maximizes power and with the default regularization parameter value of 0.05. The resulting source PSD estimates were then averaged in the delta (0.5–4 Hz), theta (4–8 Hz), alpha (8–13 Hz), beta (13–30 Hz), and gamma (40–90 Hz) frequency bands. For visualization purposes, we computed z-scores of the band-limited PSD estimates across frequency bands.

### Corticokinematic coherence analysis

We analyzed corticokinematic coherence (CKC) both at the sensor level and in the anatomical source space by computing it between the L2-norm of the 3-axis ACC signal and the preprocessed MEG epochs. For sensor-level CKC, each pair of orthogonal planar gradiometer signals was combined into a virtual gradiometer using 100 uniformly spaced angles between 0 and 180 degrees, and the angle corresponding to the strongest CKC was retained. Coherence between the ACC signal and the virtual gradiometers was estimated using the FFT coefficients in each overlapping epoch and averaged across epochs to obtain an average CKC spectrum. For the group-level analysis of CKC topographies, we used the peak movement frequency for each subject and condition, obtained from the ACC L2-norm.

To estimate CKC in source space, we applied the DICS method, which allows for straightforward computation of coherence between source-level MEG signals and an external reference signal (Gross et al. 2001). We utilized the DICS implementation in the Conpy Python package (Vliet et al. 2018). The beamformer weights were computed in the same way as for the spectral analysis above, except that separate spatial filters were obtained for each Cartesian direction in the head coordinate frame at each source point. To find the source-current orientation that maximizes coherence, we first used singular value decomposition to find the two principal source-current directions that contribute most to the MEG forward model. The optimal source-current direction at each location was then identified through a grid search across 1000 linear combinations of the principal directions. The resulting CKC maps were averaged in the 0.5–1.5-Hz range to account for variability in the movement frequency.

### Statistical analysis

We applied nonparametric permutation statistics throughout the study. From the behavioral data, we assessed the difference in the mean phase angles of the ACC phase-locking vectors across conditions using an exact two-tailed permutation t-test (8191 permutations). To investigate PSD and CKC differences between leaders and followers, we employed the two-tailed cluster-based permutation test (10000 permutations; initial cluster forming threshold *t* = 3.0) both at the sensor level and in source space (Maris and Oostenveld 2007).

## Results

### Behavioral results

Figures 1C–E summarize the behavioral results. In Figure 1C, the blue and purple traces depict the finger kinematics of the subjects at Site 1 and Site 2, respectively, illustrating how the acceleration of the follower’s finger lags that of the leader. In the top panel, Site 1 leads the movement and, hence, the purple trace lags the blue one. In the bottom panel, the roles are reversed, leading to a reversed lag. Movements were closely aligned between participants. To further investigate the behavioral dynamics, we estimated the power-spectral density (PSD) of the accelerometer signals (see Figure 1D). The grand average accelerometer (PC1) spectra attained the maximum at 0.25 Hz, but without a distinct peak, indicating that the movement frequency varied throughout the recording and across subjects.

To examine whether participants performed the task as instructed, we evaluated behavioral synchrony and the corresponding lags during leader–follower coordination as follows: We computed the phase-locking vector (PLVec) and phase-locking value (PLV) between the kinematics of the two members of each dyad separately in the conditions where participants at each site led the movement. We observed clear behavioral synchrony, with the across-subjects grand-average PLV of 0.62 when Site 1 led the movement and 0.66 when Site 2 led the movement (Figure 1E). The mean phase difference Δ*φ* between the sites was 32^∘^when Site 1 led the movement and −27^∘^when Site 2 led the movement. The observed 59 ^∘^ difference between the conditions was statistically significant (exact permutation t-test, two-tailed, 8191 permutations, *p* = 1.220 ⋅ 10^−*^, *T* = 9.272), indicating that the participants successfully followed the assigned task roles.

### Spectral signatures of leader–follower interaction

To investigate how oscillatory brain activity might support coordination between leader and follower roles, we estimated source-level power spectral density (PSD) in the delta, theta, alpha, beta, and gamma frequency bands (Figure 2). Statistically significant differences existed between leaders and followers in the theta, alpha, and beta bands (Figure 2C; cluster-based permutation test, two-sided, *p*_cluster_ < 0.05): Followers exhibited stronger theta power in a cluster spanning bilateral sensorimotor, medial frontal, and cingulate regions, including primary somatosensory cortex, primary motor cortex, premotor cortex, supplementary motor area (SMA), dorsolateral prefrontal cortex, and the mid/posterior cingulate cortex, as well as right inferior frontal regions and orbitofrontal cortex. In the alpha band, leaders showed reduced alpha power compared with followers in a cluster spanning right-lateralized sensorimotor and premotor regions, including the primary motor cortex (precentral gyrus), primary somatosensory cortex (postcentral gyrus), and SMA. In the beta-band, a significant cluster showing stronger power in leaders than followers was located in occipital regions of each hemisphere, including the lingual gyrus, pericalcarine cortex, and portions of the cuneus, as well as the right lateral occipital cortex.

**Figure 2.**
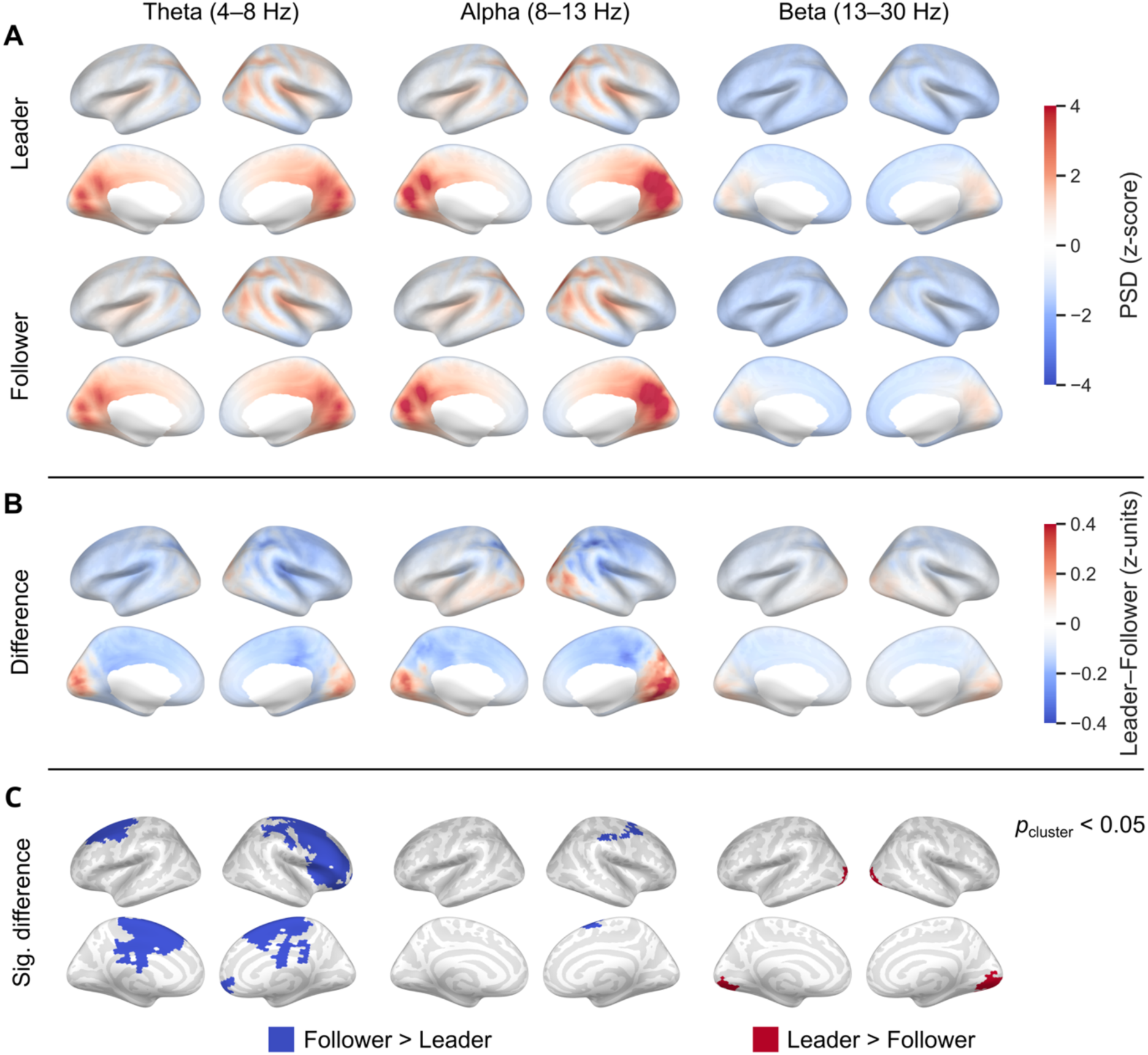
Spectral signatures of leader–follower coordination. **(A)** Grand-average source power-spectral density (PSD) in leaders and followers in the theta (4–8 Hz), alpha (8–13 Hz), and beta (13–30 Hz) frequency bands. For visualization purposes, the PSD estimates have been z-scored across both conditions and all frequency bands. **(B)** Difference in PSD between leaders and followers illustrated in z-units. **(C)** Statistically significant PSD differences between leaders and followers (cluster permutation test, two-sided).

### Brain–action coupling in the mirror game

To investigate how leaders and followers integrate visual and proprioceptive information, we estimated coupling between the brain activity of the participants and their finger kinematics (acceleration), separately for each role. Both leaders and followers showed clear corticokinematic coherence (CKC) peaks in gradiometers over bilateral sensorimotor cortices, with mean peak values of 0.3 around 1 Hz (Figure 3A). During the peak CKC, the analysis with a DICS beamformer indicated activity in the right-hemisphere sensory (S1) and motor (M1) hand areas (Figure 3A). Moreover, unlike leaders, followers also exhibited a coherence peak in occipital gradiometers, with MEG sources in visual cortical regions, predominantly in the right hemisphere.

**Figure 3.**
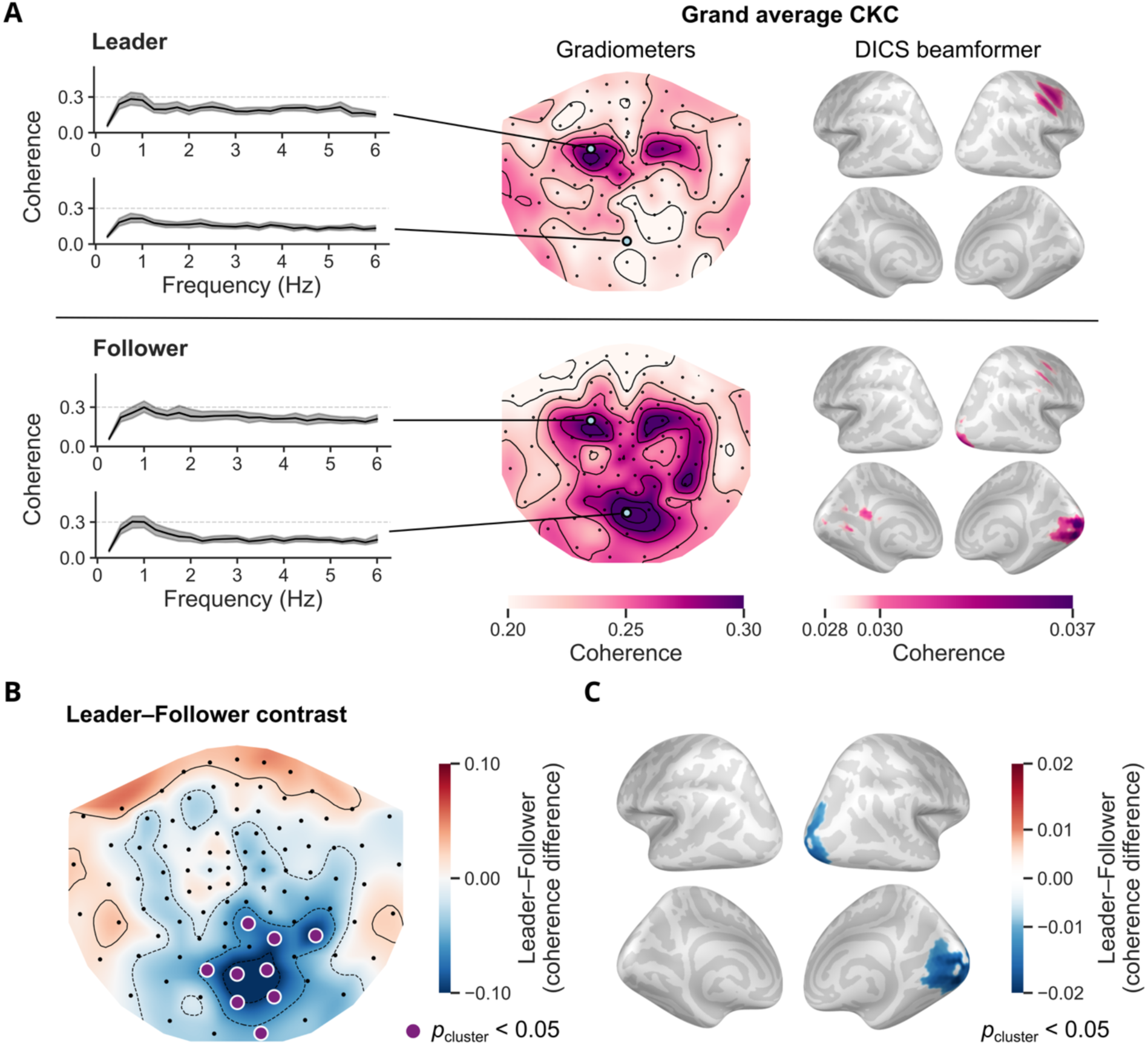
Brain–action coupling during leader–follower coordination. **(A)** Corticokinematic coherence (CKC) in leaders and followers. **Left**: Grand-average CKC spectra at selected virtual gradiometers. Shaded areas indicate the 95% bootstrap confidence interval of the mean. **Middle**: Grand average CKC virtual gradiometer topographies at the individual peak movement frequencies (the flattened map is viewed from above, with the nose up). **Right**: Grand-average DICS beamformer CKC source estimates. CKC has been averaged in the 0.5–1.5-Hz range. **(B)** Sensor-level CKC difference between leaders and followers. The significant cluster (cluster permutation test, two-sided, *p* < 0.05) is indicated with purple markers. **(C)** Source-level CKC difference between leaders and followers. Only the significant cluster (cluster permutation test, two-sided, *p* < 0.05) is displayed.

To quantify the difference between leaders and followers, we computed the leader–follower CKC contrast and assessed its statistical significance at the group level (Figure 3B). Followers showed significantly stronger CKC in a cluster of occipital gradiometers (cluster-based permutation test, two-tailed, 10 000 permutations, *p*_cluster_ < 0.05). At the source level, the same approach revealed a similar effect in a cluster in occipital regions in the right hemisphere, including the lingual gyrus, pericalcarine cortex, and portions of the cuneus. We observed no differences in sensorimotor CKC between leaders and followers. Together, these findings suggest that both leaders’ and followers’ sensorimotor areas synchronize with the movement during motor coordination, but the movement kinematics additionally entrain activity in the visual cortex of the follower.

## Discussion

In this study, we investigated the association of brain oscillatory dynamics and brain–action coupling with leader–follower coordination during mirror-game improvisation. Our findings highlight complementary patterns associated with the leader and follower roles, consistent with enhanced monitoring and adaptation to externally generated actions in followers, and internally driven action generation, motor improvisation, and top-down predictive processes in the leader role.

### Theta band

In the theta band, our results revealed a significant cluster of brain regions characterized by stronger power in followers than leaders, spanning bilateral sensorimotor, frontal, and cingulate regions. Frontal and medial theta-range activity have been consistently linked to executive control, performance monitoring, and conflict/error processing, also during interpersonal coordination (Cohen and Donner 2013; Cavanagh and Frank 2014; Ullsperger et al. 2014; Moreau et al. 2020; Boukarras et al. 2022; Cavanagh and Cohen 2022; Pyasik et al. 2023). Frontocentral theta has been shown to increase when the subject coordinates with a virtual partner that suddenly and unpredictably changes its movement trajectory, violating action predictions (Moreau et al. 2020). Theta-range activity has even been suggested to have a causal role in facilitating action monitoring and interpersonal motor coordination (Boukarras et al. 2022). Complementary evidence shows that followers exhibit stronger frontocentral error-related responses to unexpected corrections performed by an interaction partner than to their own subsequent adaptations (Moreau et al. 2023), as well as stronger neural tracking of visual self–other synchronization errors during mirror-game interaction (Varlet et al. 2025). Considered together with this literature, the stronger theta-band activity we observed in the follower role may reflect heightened demands for monitoring (self- and other-generated) actions and coordination errors to sustain sensorimotor coordination under the uncertainty of motor improvisation.

### Alpha band

We observed a set of brain regions with weaker alpha power in leaders than followers, covering right-lateralized sensorimotor and premotor areas. A reduction in alpha power is generally interpreted to reflect increased cortical excitability and selective processing of task-relevant information, supporting the gating and routing of information toward task-relevant regions while inhibiting competing or task-irrelevant processes (Klimesch et al. 2007; Jensen and Mazaheri 2010; for a review, see Jensen and Bonnefond 2026). Given that leading in the mirror game requires producing novel movement trajectories, the Rolandic alpha suppression observed in leaders may reflect greater recruitment of motor-oriented networks supporting self-generated action planning and motor improvisation processes.

This interpretation is consistent with previous fMRI studies showing stronger recruitment of premotor and motor-related regions during leading in joint movement improvisation tasks (Fairhurst et al. 2014; Chauvigné and Brown 2018). The site of our significant cluster in right sensorimotor and premotor regions (Figure 2C, middle panel) overlaps with cortical regions previously associated with motor improvisation and expertise in the leader role (Beaty 2015; Chauvigné and Brown 2018). Beyond movement-related activity, Konvalinka and colleagues (2014) reported stronger suppression of EEG alpha activity in frontal regions in spontaneously emerging leaders during synchronized finger tapping guided by auditory cues, and they interpreted it to indicate increased prospective planning and control. However, given its frontal distribution and lack of source analysis in the recordings with a nasal reference, this effect likely reflects a different alpha generator from the Rolandic sensorimotor alpha activity implicated in the present findings. Nonetheless, taken together, our findings converge with this literature to suggest that leading relies more strongly on cortical activity supporting internally driven action planning and control.

It is worth noting that, since both participants performed movements with the right hand, the right-lateralized distribution of the Rolandic alpha effect is unlikely to be explained by low-level movement execution processes, which would be expected to produce predominantly contralateral or bilateral centroparietal alpha-band suppression. Instead, the findings are broadly consistent with previous reports of right-lateralized centroparietal EEG alpha/mu activity during interpersonal coordination (Naeem et al. 2012b, 2012a; Novembre et al. 2016). Importantly, given that these studies utilized EEG and reported predominantly centroparietal effects, it is possible that some of their findings were influenced by parietooccipital alpha contributions, which differ functionally from the more central activity observed in the present study. Compared with EEG, MEG offers better separability of simultaneously active current sources (Hämäläinen et al., 1993; Hari and Salmelin, 2012). The present findings extend this literature by identifying role-dependent right-lateralized Rolandic alpha-band differences during joint improvisation.

### Beta band

In the beta band, we identified a significant cluster of bilateral occipital areas with increased power in leaders compared with followers. Beta oscillations have been implicated, among others, in endogenous top-down signaling, feedback and predictive processes, and in the maintenance of ongoing cognitive and sensorimotor states (Engel and Fries 2010; Cannon et al., 2014; Bastos et al. 2015; Jensen et al. 2015; Zheng and Colgin 2015). Within this framework, beta-band activity has been proposed to convey moment-to-moment task demands and behavioral context to sensory regions, including the visual cortex (Richter et al. 2018). Such top-down influences have been shown to shape early visual processing according to current behavioral demands (Jensen et al. 2015; Richter et al. 2018; Roehe et al. 2023). Moreover, action planning and motor preparation influence visual cortical activity, even in the absence of visual feedback, and can predictively shape visual processing in preparation for sensory changes contingent on self-generated action (Astafiev et al. 2004; Orlov et al. 2010; Kühn et al. 2011; Lingnau and Downing 2015; Benedetto et al. 2021). In the present task, leaders determined the upcoming movement directions, amplitudes, and temporal structure and could therefore predict a substantial part of their visual input, while followers had to rely on the received visual information. We propose that the enhanced occipital beta activity observed in leaders reflects stronger top-down predictive influences on visual processing, shaped by internally generated motor plans and expectations regarding the anticipated sensory consequences of their actions.

This interpretation complements recent findings from a mirror-game setup showing role-dependent differences in the neural tracking of self–other synchronization errors (Varlet et al. 2025). Leaders exhibited similar latencies of error tracking across motor and visual areas, while followers showed delays, potentially reflecting the time required for error-related information to travel bottom-up and be processed and integrated into ongoing behavior. Varlet and colleagues (2025) interpreted this pattern as evidence for stronger predictive processing and top-down influences from motor to visual systems when leading. Accordingly, the stronger occipital beta activity observed in leaders in the present study may reflect a related mechanism through which improvised motor plans exert greater top-down influence on visual processing.

Notably, a previous study by Zhou and colleagues (2016), examining leader–follower coordination during hand opening-and-closing movements, did not show significant differences in alpha- or beta-band power between roles (Zhou et al. 2016). However, the work involved highly repetitive and thus predictable movements, whereas the present mirror-game task relied on movement improvisation and thus yielded less predictable movement patterns. The alpha- and beta-band differences observed here may therefore reflect neural processes that become particularly relevant during joint improvisation, such as internally driven prospective planning and top-down predictive control in the leader role.

### Brain–action coupling

Our CKC analysis revealed a significant role-dependent difference, with stronger coherence between movement kinematics and right occipital cortical activity in followers than in leaders. This occipital effect should be distinguished from sensorimotor cortico-kinematic coherence (CKC) observed in both roles (Jerbi et al. 2007; Bourguignon et al. 2011). The classic sensorimotor CKC is thought to primarily reflect proprioceptive afferent input to the cortex (Piitulainen et al., 2013; Bourguignon et al., 2015). Related studies have also reported neural tracking of observed actions beyond sensorimotor regions, with the kinematics of observed movements coupled to activity in visual, parietal, and cerebellar regions (Bourguignon et al., 2013; Marty et al., 2015). The stronger occipital CKC observed in followers may therefore reflect enhanced visual tracking of movement kinematics rather than stronger proprioceptive sensorimotor coupling. This result is consistent with the task demand. Although both participants had visual access to their partner’s movements, the visual information was more relevant for the followers, who were required to continuously monitor the leader and adjust their own movements accordingly.

Our results suggest that leaders and followers process the available visual kinematic information differently, consistent with prior evidence that followers rely more on visual feedback and self–other integration processes for movement control and coordination (Zhou et al. 2016; Varlet et al. 2020, 2025). Previous frequency-tagging studies have shown stronger neural responses to the partner’s movements in followers, while leaders exhibited enhanced tracking of self-generated motion (Varlet et al. 2020). In line with this literature, the right-lateralized occipital distribution of our CKC difference is compatible with stronger tracking of information presented in the contralateral (left) visual hemifield, corresponding to the location of the leader’s hand movements on the screen. Our findings also align with the evidence for enhanced neural tracking of visual self–other synchronization errors in followers compared with leaders, which have been interpreted to reflect greater adaptation demands and stronger reliance on incoming sensory evidence during coordination (Varlet et al. 2025). Moreover, our results converge with previous evidence that the kinematic phase of hand movements modulates occipital beta-band activity more strongly in followers than in leaders, which may be indicative of distinct role-dependent strategies for integrating visual kinematic information during interaction (Zhou et al. 2016). The present findings extend this literature by providing evidence that direct, frequency-specific brain–action coupling is modulated by leader–follower roles during interpersonal coordination.

### Limitations and future directions

A few methodological considerations should be recognized when interpreting our results. Although statistically significant role-dependent effects were identified using cluster-based permutation testing, our sample size and recording duration were modest. Replication in larger samples and longer recordings will be important to establish the robustness and generalizability of the reported findings. We should also note an intrinsic limitation of cluster-based statistics: statistical significance was established at the cluster level, indicating a significant difference somewhere within the cluster, but not implying that a significant change occurs at every individual brain region or sensor within the cluster (Maris and Oostenveld 2007; Sassenhagen and Draschkow, 2019).

Second, CKC is typically strongest for rhythmic movements with stable movement frequencies, whereas our mirror-game task required continuous movement improvisation. Therefore, movement frequencies varied across participant pairs and over time during the interaction, attenuating coherence estimates. Although our analyses were tailored to individual movement frequencies derived from accelerometer data, the effect sizes were (expectedly) smaller than those reported in highly rhythmic paradigms (Bourguignon et al. 2011; Piitulainen et al. 2013). Nonetheless, CKC provides a valuable approach for examining continuous brain–action coupling during interpersonal interaction. Future hyperscanning studies should integrate brain–action measures such as CKC with interbrain analysis and thus provide additional insights into the brain basis of real-world interpersonal coordination (Hari et al. 2015; Avendano-Diaz et al., 2026).

More broadly, the present findings reflect a specific mode of coordination and should not be assumed to generalize to all forms of leader–follower dynamics. Previous studies have reported distinct oscillatory signatures across varying task demands, sensory modalities, and interaction structures (Konvalinka et al. 2014; Zhou et al. 2016; Yoneta et al. 2022). Factors such as visual versus auditory coordination, spontaneous versus predefined leadership roles, ongoing versus asynchronous coordination, or direct versus indirect access to a partner’s behavior may influence the neural processes manifesting leader–follower dynamics. The present findings are likely most representative of continuous visuomotor joint improvisation with predefined roles and direct visual access to a partner’s behavior. Further work could investigate whether universal spectral signatures of leader–follower coordination emerge across diverse forms of interpersonal interaction. In addition, future studies spanning a continuum from tightly controlled to more naturalistic task designs may help clarify whether and to which extent the different spectral signatures reported here might have specific functional roles in interpersonal coordination.

Overall, the present findings extend previous neuroimaging work on movement interaction by identifying spectral and brain–action coupling signatures associated with leader–follower dynamics during joint improvisation, offering new insights into the neural basis of real-time interpersonal coordination.

## Author contributions statement

According to the Contributor Role Taxonomy (CRediT), the authors contributed to the study as follows: Conceptualization, Methodology, Project administration, Writing – review & editing – S.R., J.C.A.-D., R.H., L.P.; Data curation, Formal analysis, Investigation, Validation, Writing – original draft – S.R., J.C.A.-D.; Software, Visualization - S.R.; Funding acquisition – J.C.A.-D., R.H., L.P.; Supervision, Resources – R.H, L.P.

## Acknowledgments

We thank Pamela Baess, Marc Op de Beeck, Mathieu Bourguignon, Matti Hämäläinen, Tommi Himberg, Lotta Hirvenkari, Mia Illman, Veikko Jousmäki, Mathilde Menoret, Jyrki Mäkelä, Anne Mandel, Jussi Nurminen, Petteri Räisänen, Ronny Schreiber, Sara Sommariva, Arianna Schiano, Teemu Taivainen, Marijn van Vliet, Andrey Zhdanov, and Xavier de Tiège at Aalto University and Universitè Libre de Bruxelles for their contributions to various stages of data collection and analysis. We thank Matti Hämäläinen for providing resources and insight on methodology. We acknowledge the computational resources provided by the Aalto Science-IT project. Artificial intelligence tools were used in part to create the illustrative parts of Fig 1A and Fig 1B. This work was supported by the European Research Council (Advanced Grant Brain2Brain, award number 232946 to Riitta Hari); the National Institute of Neurological Disorders and Stroke (award number 7R01NS104585-06 to Matti Hämäläinen); Business Finland (grant “DIGIMIND” 7981/31/2022); The Norman Loveless Memorial Fund; and the Ella C Georg Ehrnrooth Foundation.

